# GenesetDiseaseDrugNetwork (GDDN): a web server for disease enrichment and drug prioritization

**DOI:** 10.64898/2026.05.22.727145

**Authors:** Piyush More, Jean-Fred Fontaine, Vincent ten Cate, Philipp S. Wild, Miguel A. Andrade-Navarro

## Abstract

**Summary:** Omics technologies profile thousands of genetic and molecular features to provide a comprehensive and quantitative measure of the cellular state. Transcriptomics and proteomics have, especially, guided discoveries of the most important biomarkers and therapeutic targets. By virtue of ongoing developments in single-cell and spatial technologies, fields of targeted therapeutics and personalized medicine are rapidly advancing. However, downstream functional analysis and disease association still remain daunting tasks in bioinformatics. We address these challenges with the GenesetDiseaseDrugNetwork (GDDN) web server. GDDN facilitates functional discovery by connecting gene-sets to enriched diseases and their corresponding therapeutics in a single step. Using a ranking system that incorporates regulatory impact, specificity, and potency, GDDN effectively prioritizes drugs with the highest clinical relevance. Our platform facilitates the interpretation of omics outputs into disease associations and personalized drug identification.

**Availability and Implementation:** The GDDN web server is implemented in R Shiny and is freely accessible at https://cbdm-01.zdv.uni-mainz.de/shiny/piyusmor/GDDN/, supporting all major web browsers.

**Supplementary information:** Supplementary data is available at Bioinformatics online.

## 1 Introduction

Omics and next generation sequencing platforms have transformed our understanding of biology. Genomics, transcriptomics, epigenomics, and proteomics provide crucial information regarding genome, gene expression, regulation, and protein dynamics even at a single cell resolution. This has radically revolutionized genetic research and clinical applications. Whole genome or targeted sequencing is routinely performed in cancer diagnostics to guide specific treatments. BRCA1/2 screening in breast cancers, EGFR mutations in lung cancer, PD-L1 profiling for immunotherapy, and troponin for diagnosing myocardial infarction are some of the most successful clinical examples (Haaf *et al*. 2012, Rituraj *et al*. 2025).

Despite the clinical success, post-quantification downstream analysis of omics data remains challenging. Two of the most pivotal challenges in biomedical research are interpreting the biological context of large number of target-sets and deriving effective therapeutic intervention strategies. There are few methods and tools that are widely used to independently address above mentioned tasks. The Open Targets Platform (OTP) provides a comprehensive suite for target-disease association and drug prioritization (Buniello et al. 2025). However, OTP considers unique target-disease pairs; thereby, the discovery pipeline focuses on individual associations rather than the combined contributions of multiple genes. Analogously, DisGeNET also focuses on individual gene-disease pairs (Piñero et al. 2026). GDIdb is a database used to identify drugs potentially modulating set of genes based on known gene-drug interactions and predicted “druggable” genes (Cannon et al. 2024). There are databases like DrugCentral (Ursu et al. 2017), DrugBank (Knox et al. 2024), and ChEMBL (Bento et al. 2014) for connecting diseases or gene targets to drugs. A systematic method for associating gene sets (as derived from various omics technologies), first, to diseases and then to drugs is therefore missing.

To fill this void, we developed the GenesetDiseaseDrugNetwork (GDDN) resource. GDDN provides an integrated method of statistically associating functionally related gene sets to diseases and identifying and raking potential drugs candidates. The GDDN web server is built on publicly available knowledge. It is based on gene and disease annotations in biomedical records and validated information regarding drug-target interactions and drug indications. We envision that GDDN will enable the functional interpretation of omics-derived gene and protein sets and will uncover potential drug candidates, by leveraging those gene-disease associations.

## 2 Methods

### 2.1 Datasets and web-server

We obtained all the datasets used in this work from public repositories. First, we retrieved year 2025 annual baseline and daily update files from PubMed FTP servers (https://pubmed.ncbi.nlm.nih.gov/download/). We then obtained genes to PubMed mappings (genes2pubmed.gz) from the NLM website (https://ftp.ncbi.nlm.nih.gov/gene/DATA/). We obtained medical subject headings from the Medical Subject Headings (MeSH) database (branch C for disease terms). Finally, we processed the DrugBank database (Knox et al. 2024) to extract drugs, nomenclature, targets, and clinical indications. We built the GDDN web-server using the Shiny package in R (Chang et al. 2025). To enable JavaScript functionality, we utilized the shinyjs package in R (Attali 2026).

### 2.2 Gene-disease association

GDDN establishes gene-disease associations by cross-referencing gene and clinical annotations within the biomedical literature (PubMed). Using the genes2pubmed.gz file, we identified all PubMed IDs (PMIDs) linked to human or mouse genes and stored them in a MySQL database restricted to records that are simultaneously annotated with at least one gene and one MeSH C disease term. For comparison between human and mouse orthologs, ENSEMBL database annotations were used (Yates et al. 2022). We then derived significant associations of gene and disease using the one-tailed Fisher’s exact test (Fontaine and Andrade-Navarro 2016, More et al. 2021). For each gene-disease association, we derived a 2×2 contingency matrix consisting of following four groups -PMIDs annotated with both gene and disease, PMIDs annotated with only the gene and not the disease, PMIDs annotated with only the disease and not the gene, and PMIDs annotated with neither the gene nor the disease. To correct for multiple testing, we calculated the false discovery rate (FDR) by the Benjamini and Hochberg method in R (Benjamini and Hochberg 1995, R Core Team 2025). We then derived significant gene-disease associations by identifying co-occurrences with at least three PMIDs and less than 0.05 FDR. These curated associations are used for disease enrichment analysis.

GDDN can be started with a user’s submitted gene set. Disease-enrichment analysis is performed using a one-tailed Fisher exact test on this input gene-set. To achieve this, for each disease, we derive a contingency matrix of number of genes from the submitted gene-set associated with the disease, number of genes from the background gene-set associated with the disease, number of genes from the submitted gene-set not associated with the disease, and number of genes from the background gene-set not associated with the disease. Analogous to the previous step, we calculated the FDR using the Hochberg and Benjamini method.

### 2.3 Drug prioritization

To allow the identification of top drug candidates, we utilized the information regarding drug targets and drug indications. We extracted nomenclature, clinical indications, and targets from the DrugBank database (Knox et al. 2024). To normalize DrugBank’s descriptive indications into formal disease terms, we utilized a local BERTMesh LLM model specifically trained on the MeSH thesaurus (You et al. 2021). Resulting terms were further refined using the MeSH lookup API. This step was essential to derive connections between PubMed records and DrugBank entries.

Given a set of diseases found to be enriched in a submitted gene set in the previous step, we next identify drugs indicated for those diseases as well as drugs targeting the genes within the set. For drug prioritization, we calculated relevance, regulatory impact, specificity, and binding affinity for each drug. To derive relevance, we count the number of proteins in the input gene-set, targeted by the drug. To derive regulatory impact, we investigate whether the drug targets transcription factors, as defined in the Transcription Factors database, (Lambert et al. 2018) or other proteins involved in key cellular regulatory processes. Those processes include DNA-binding transcription factor activity (GO:0003700), chromatin remodeling (GO:0016590), and protein kinase activity (GO:0004672). We then assign the regulatory score (1 or 0) to either presence or absence of transcriptional factors or regulatory processes. We use a specificity score as 1/n (where n is the number of targets). To derive binding affinity, we first obtained pChEMBL scores for each drug from the ChEMBL database (Bento et al. 2014). If the pChEMBL score was not available, we assigned a mean score, calculated across all drugs. We then normalized all four scores, relevance_score, regulatory_impact_score, specificity_score, and binding_affinity_score, between 0 and 1. We finally calculate the final drug_priority_score by the weighted sum of all four scores. For this, we assign 50% weight to relevence_score, 30% weight to regulatory_impact_score, 15% weight to specificity_score, and 5% weight to binding_affinity_score.

## 3 Results

### 3.1 Data overview

By associating genes to diseases, we can understand the biological context of a particular cellular state. We can also postulate the roles of genes within a specific disease or across multiple diseases, helping us to decipher the associated regulatory networks.

Disease enrichment analysis is performed on a set of genes and relies on filtered biomedical annotations in the literature. We have identified statistically significant co-occurrences of genes and diseases in the PubMed and MeSH databases (see Methods). This forms the basis of the GDDN web-server. GDDN contains a total of 873,299 associations between 43,946 genes and 4321 diseases. Among those, there are 78,625 associations between 12,678 genes and 2692 diseases having more than three PubMed citations and FDR < 0.05. Considering that substantial evidence for gene involvement in human diseases stems from *in vivo* mouse models, we extended our approach to include mouse genes, leveraging orthology information to facilitate seamless integration. Among 78,625 significant gene-disease associations, 75,813 are in humans (11869 genes and 2616 diseases) and 2812 are in mice (809 genes and 706 diseases).

### 3.2 Web server implementation

The interactive web-server provides three analysis modes, Gene-set enrichment, Genes to Diseases, and Diseases to Genes. Gene-set enrichment first performs disease enrichment analysis on a set of genes, where the association of multiple genes is considered simultaneously. Then identifies and ranks drugs based on shared targets and enriched diseases. Figure 1 shows the overview of the web server implementation. Users can select the number of genes to be associated with a particular disease (minimum two, default two) and number of PubMed citations supporting it (minimum two, default five) to be considered for the analysis. The tabular output provides enriched diseases ranked according to FDR and the number of genes. The table also provides external links mapping genes to NCBI Entrez IDs and Ensembl IDs, and PMIDs to respective PubMed articles. The enrichment results are also summarized in the form of an UpSet plot. It shows the top genes shared by enriched diseases and facilitates the identification of related diseases in terms of common genes. The web-server also provides a Drug Prioritization table with a list of drugs (ranked as explained in Methods) and their gene targets including external links to DrugBank online and NCBI or Ensembl databases. To visualize the connection between drugs and genes, an interaction network is also presented. To avoid large networks, this is generated for individual diseases. In this network, the central node in red represents the disease. The nodes in orange and blue color represent genes (symbols) and drugs respectively.

**Figure 1:**
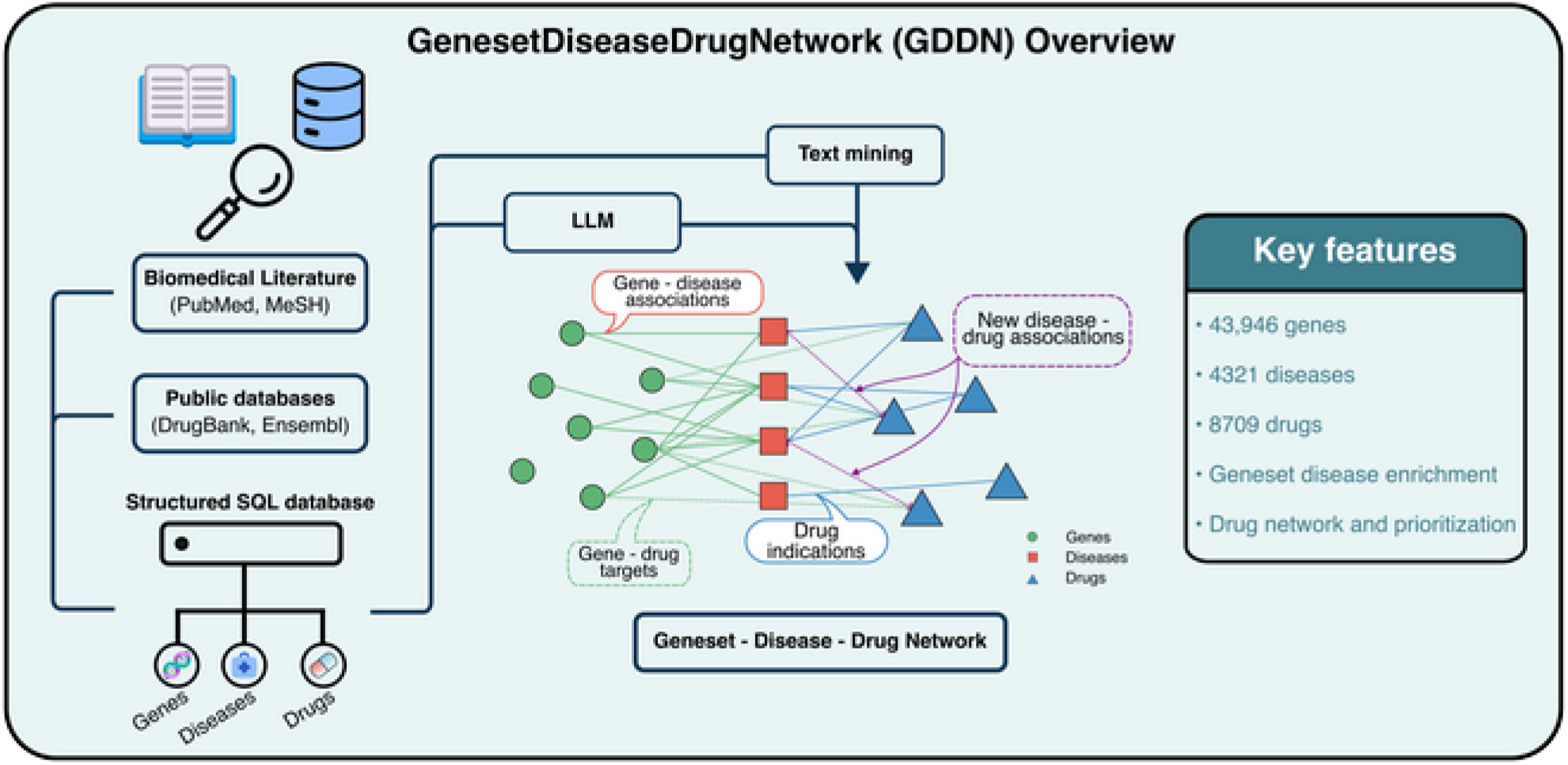
Graphical representation of the GenesetDiseaseDrugNetwork (GDDN) web server implementation.

Further analysis modes, Genes to Diseases and Diseases to Genes provide associations of individual genes and individual diseases respectively. Those associations are derived from pre-computed significant co-occurrences of genes and diseases (see Methods for details). Users can modify the minimum number of PubMed citations supporting the gene-disease association (minimum two, default five) and the FDR threshold (default 0.05) to adjust the stringency of results. All results can be downloaded locally as tabular data in TSV format and graphical results as PDF or PNG image files. The users can use pre-defined examples to understand the input format and to test the functionality of the web-server.

### 3.3 Use case

To demonstrate the utility of the GDDN web-server, we identified 302 genes overexpressed in TP53-mutated liver hepatocellular carcinoma (LHPC) using the cBioportal platform (Supplementary Table 1). There is a total of 115 patient samples with TP53 mutations (90 males and 25 females) and 258 samples without TP53 mutation (162 males and 96 females) in the Liver Hepatocellular carcinoma TCGA Firehose Legacy cohort. The clinical data shows worse overall survival in the mutated group (mean overall survival 45 months) compared to the non-mutated group (mean overall survival 61 months) (Supplementary Figure 1). Using the GDDN web-server, we identified 52 diseases significantly enriched within the set of 302 genes. Among those diseases, hepatocellular carcinoma was associated with 50 genes followed by liver neoplasms, squamous cell carcinoma, lung neoplasms, and genome instability (Supplementary Figure 2 and Supplementary Table 2). The enrichment plot showed related diseases based on shared genes. These included hepatocellular carcinoma and liver neoplasms, chromosome instability and aneuploidy, and melanoma and synovial sarcoma (Supplementary Figure 3). The GDDN ranked potential drugs to include Fostamatinib, Clonazepam, Flumazenil, Flunitrazepam, and Brotizolam among others (Supplementary Figure 4). Fostamatinib targets MELK, NEK2, PLK1, PKMYT1, and TTK, while the remaining four drugs target GABRA2 and GABRA3. Fostamatinib is an investigational kinase inhibitor with several clinical trials (NCT00923481, NCT00446095) and promising activity, especially, against hematological malignancies (Park *et al*. 2013, Hu *et al*. 2025).

## 4 Conclusion

The GDDN web server provides a unified method to associate genes with diseases and drugs. It is a unique tool performing disease enrichment on a set of genes and providing ranked lists of drugs by accounting for biological relevance. GDDN can be accessed freely using all major web browsers in a user-friendly manner. All results can be downloaded locally for further investigations.

## Acknowledgments

The authors gratefully acknowledge the help from the IT group at the Johannes Gutenberg University Mainz. They would also like to thank Celine Mueller and Markus Ingold for testing the web server and for their suggestions.

## Funding

This work was supported by funding from the Federal Ministry of Research, Technology and Space of Germany (BMFTR) Clusters4Future (C4F) initiative; curATime, cluster for Atherothrombosis and individualized medicine to M.A. (funding numbers: 03ZU1202AA and 03ZU1202EC).

## Notes

### Competing Interest Statement

The authors have declared no competing interest.

