## Supplementary Information for "GenesetDiseaseDrugNetwork (GDDN): a web server for disease enrichment and drug prioritization"

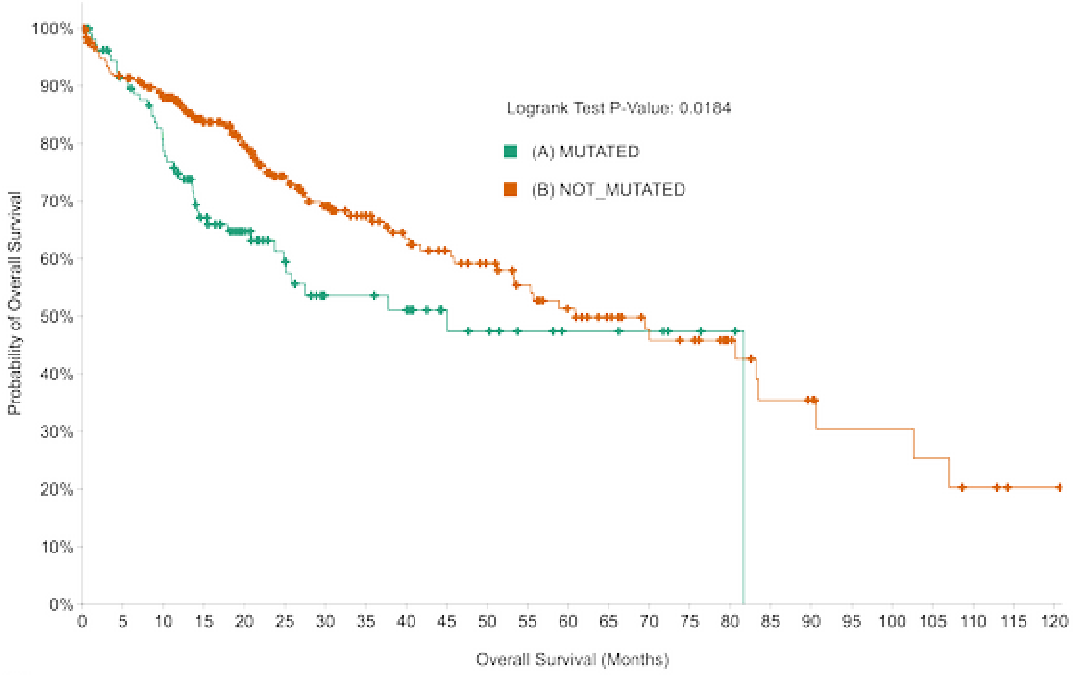


**Number at risk (n)**

| | | | | | | | | | | | | | | | | | | | | | | | | | |
|---|---|---|---|---|---|---|---|---|---|---|---|---|---|---|---|---|---|---|---|---|---|---|---|---|---|
| (A) MUTATED | 115 | 95 | 80 | 59 | 44 | 32 | 22 | 22 | 20 | 14 | 12 | 8 | 6 | 6 | 5 | 3 | 2 | 0 | 0 | 0 | 0 | 0 | 0 | 0 | 0 |
| (B) NOT_MUTATED | 257 | 222 | 204 | 164 | 136 | 106 | 87 | 75 | 62 | 55 | 49 | 42 | 35 | 29 | 24 | 22 | 15 | 10 | 9 | 6 | 6 | 5 | 3 | 1 | 1 |

Supplementary Figure 1: The Kaplan-Meier plot comparing overall survival between hepatocellular carcinoma with and without TP53 mutation.

**Enrichment Table**

| disease_mesh_id | disease_mesh_term | gene_id | gene_symbol | gene_ensembl | gene_biotype | n_gene | pmid | p_val | fdr_bh |
|---|---|---|---|---|---|---|---|---|---|
| D006528 | Carcinoma, Hepatocellular | 332 891 990 991 1033 ... (show 45 more) | BIRC5; CCNB1; CDC6; CDC20; CDKN3 ... (show 46 more) | ENSG00000089685 ENSG00000134057 ENSG00000094804 ENSG00000117399 ENSG00000100526 ... (show 57 more) | protein_coding | 50 | 559 | 6.88e-8 | 0.00000379 |
| D008113 | Liver Neoplasms | 332 891 990 991 1033 ... (show 38 more) | BIRC5; CCNB1; CDC6; CDC20; CDKN3 ... (show 38 more) | ENSG00000089685 ENSG00000134057 ENSG00000094804 ENSG00000117399 ENSG00000100526 ... (show 46 more) | protein_coding | 43 | 597 | 0.0000712 | 0.000803 |
| D002294 | Carcinoma, Squamous Cell | 332 701 768 990 1029 ... (show 26 more) | BIRC5; BUB1B; CA9; CDC6; CDKN2A ... (show 27 more) | ENSG00000089685 ENSG00000156970 ENSG00000107159 ENSG00000094804 ENSG00000147889 ... (show 36 more) | protein_coding | 31 | 842 | 1.84e-8 | 0.00000135 |
| D008175 | Lung Neoplasms | 332 1029 1485 1869 1894 ... (show 24 more) | BIRC5; CDKN2A; CTAG1B; E2F1; ECT2 ... (show 25 more) | ENSG00000089685 ENSG00000147889 ENSG00000184033 ENSG00000101412 ENSG00000114346 ... (show 33 more) | protein_coding;lncRNA | 29 | 700 | 0.00496 | 0.0273 |
| D042822 | Genomic Instability | 641 699 701 891 991 ... (show 22 more) | BLM; BUB1; BUB1B; CCNB1; CDC20 ... (show 22 more) | ENSG00000197299 ENSG00000169679 ENSG00000156970 ENSG00000134057 ENSG00000117399 ... (show 22 more) | protein_coding | 27 | 235 | 0.000180 | 0.00188 |

Supplementary Figure 2: The GDDN enrichment results showing top 5 diseases associated with 302 genes over-expressed in *TP53*-mutated hepatocelullar carcinoma.

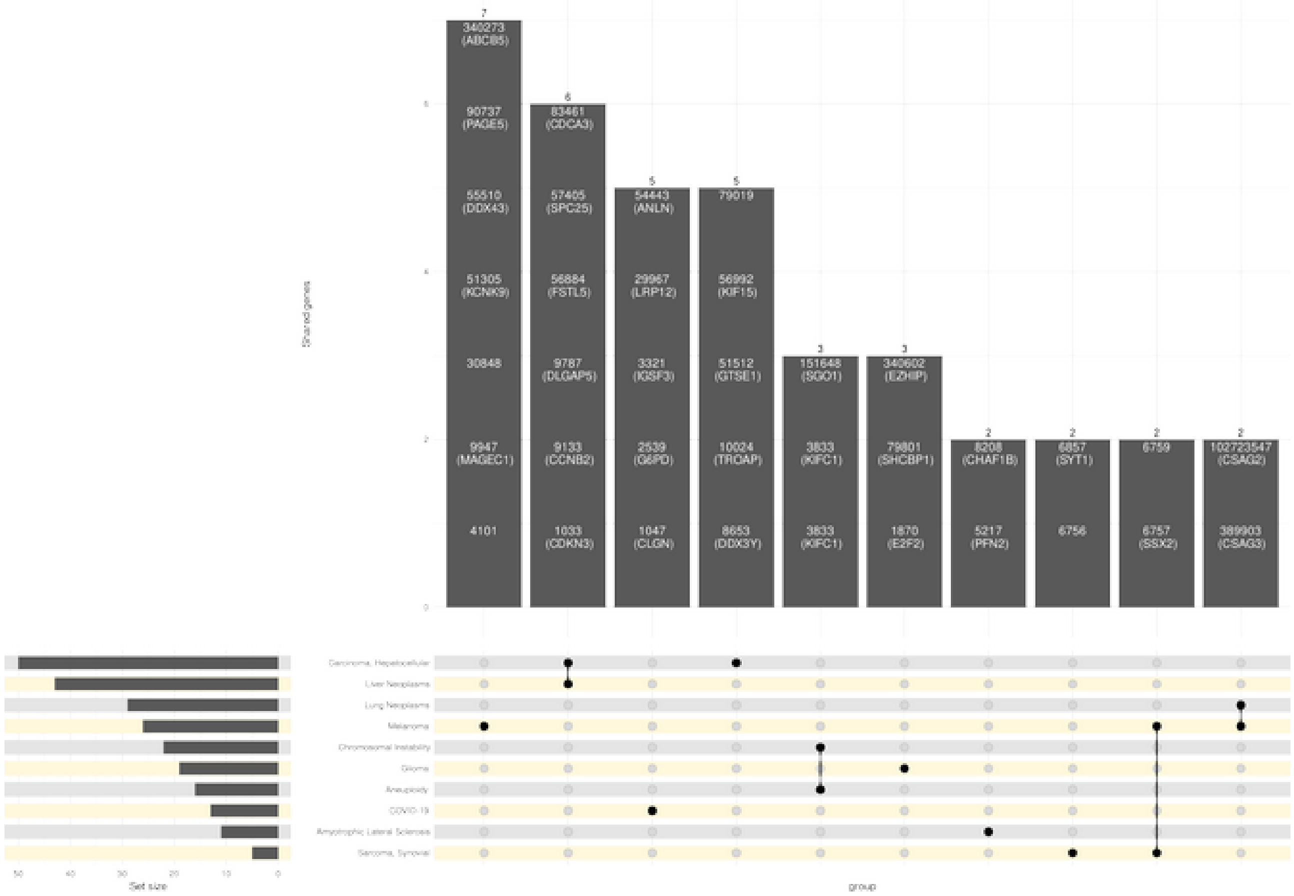


Supplementary Figure 3: The UpSet plot generated by the GDDN summarizing disease associations. The bars in the top panel show the number and ENTREZ Ids/Symbols of the genes shared between diseases. The bottom panel shows the diseases and interconnections.

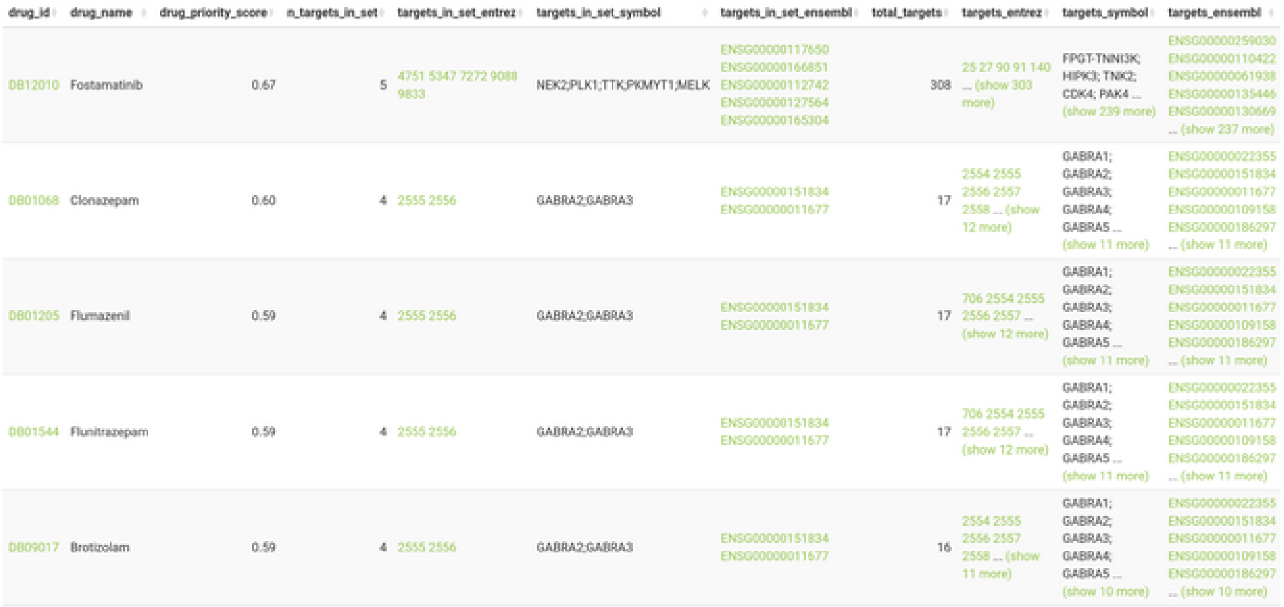

| drug_id | drug_name | drug_priority_score | n_targets_in_set | targets_in_set_entrez | targets_in_set_symbol | targets_in_set_ensembl | total_targets | targets_entrez | targets_symbol | targets_ensembl |
|---|---|---|---|---|---|---|---|---|---|---|
| DB12010 | Fostamatinib | 0.67 | 5 | 4751 5347 7272 9088 9833 | NEK2;PLK1;TTK;PKMYT1;MELK | ENSG00000117650 ENSG00000166851 ENSG00000112742 ENSG00000127564 ENSG00000165304 | 308 | 25 27 90 91 140 ... (show 303 more) | FPGT-TNNI3K; HIPK3; TNK2; CDK4; PAK4 ... (show 239 more) | ENSG00000259030 ENSG00000110422 ENSG00000061938 ENSG00000135446 ENSG00000130669 ... (show 237 more) |
| DB01068 | Clonazepam | 0.60 | 4 | 2555 2556 | GABRA2;GABRA3 | ENSG00000151834 ENSG00000011677 | 17 | 2554 2555 2556 2557 2558 ... (show 12 more) | GABRA1; GABRA2; GABRA3; GABRA4; GABRA5 ... (show 11 more) | ENSG00000022355 ENSG00000151834 ENSG00000011677 ENSG00000109158 ENSG00000186297 ... (show 11 more) |
| DB01205 | Flumazenil | 0.59 | 4 | 2555 2556 | GABRA2;GABRA3 | ENSG00000151834 ENSG00000011677 | 17 | 706 2554 2555 2556 2557 ... (show 12 more) | GABRA1; GABRA2; GABRA3; GABRA4; GABRA5 ... (show 11 more) | ENSG00000022355 ENSG00000151834 ENSG00000011677 ENSG00000109158 ENSG00000186297 ... (show 11 more) |
| DB01544 | Flunitrazepam | 0.59 | 4 | 2555 2556 | GABRA2;GABRA3 | ENSG00000151834 ENSG00000011677 | 17 | 706 2554 2555 2556 2557 ... (show 12 more) | GABRA1; GABRA2; GABRA3; GABRA4; GABRA5 ... (show 11 more) | ENSG00000022355 ENSG00000151834 ENSG00000011677 ENSG00000109158 ENSG00000186297 ... (show 11 more) |
| DB09017 | Brotizolam | 0.59 | 4 | 2555 2556 | GABRA2;GABRA3 | ENSG00000151834 ENSG00000011677 | 16 | 2554 2555 2556 2557 2558 ... (show 11 more) | GABRA1; GABRA2; GABRA3; GABRA4; GABRA5 ... (show 10 more) | ENSG00000022355 ENSG00000151834 ENSG00000011677 ENSG00000109158 ENSG00000186297 ... (show 10 more) |

Supplementary Figure 4: The drug prioritization table generated by the GDDN. It shows the top five ranked drugs potentially targeting genes over-expressed in *TP53*-mutated hepatocellular carcinoma.
